# Studies on effect of gold nanoparticles on *Meloidogyne incognita* and tomato plants growth and development

**DOI:** 10.1101/428144

**Authors:** Rajni Kant Thakur, Babita Dhirta, Poonam Shirkot

## Abstract

The plant parasitic nematodes are one of world major agricultural pest, causing in excess of 157 billion dollars in worldwide damage annually. This study has provided evidence that gold nanoparticles have great utility for management of root-knot nematodes in tomato crop. The effect of gold nanoparticles on *Meloidogyne incognita* J2 was remarkable under the direct exposure in water, after three hours of incubation of Meloidogyne incognita with GNPs showed the 100% mortality. The lesser survival rate of *Meloidogyne incognita* in soil treatment showed the strong nematicidal effect of gold nanoparticles. Subsequently, the pot experiment had shown the beneficial effects of gold nanoparticles for intensively managing the root-knot nematode. The Pot experiment not only showed us that GNPs were lethal to root-knot nematodes were also induces growth of tomato plants and didn’t have any kind of negative impact on plant growth. In our study, GNPs were found to be safe and lethal to Meloidogyne incognita.

## Introduction

The bio-gold nanoparticles synthesized by microbes are the result of various molecular strategies to overcome the mental stress by decreasing redox state of metal electron shuttle which can be extracellular or intracellular, leading to the conversion of gold metal ions to nanoparticles of defined shape and size (Konishi et al., 2006). The biosynthetic approach is particularly significant over the chemical synthesis leading to generation of toxic substances and expenditure of heavy metal and the nanoparticles have a tendency to clump together and are rendered useless nanoparticles produced by physical method are unstable and tend to agglomerate (Ali et al., 2012, Saldan et al., 2012). The advantage of nanoparticles synthesized by various living systems is a coating with peptide or protein which tends to uniform charge distribution all over the surface of nano-metal resulting in repulsion between them (Palomo et al., 2016). Thus gold nanoparticles are extremely stable even after months. Bacteria have the utmost ability to pop in metal ions and conglomerate within the cell without any harm. The bio-gold nanoparticles were first synthesized by (Beveridge and Murry in 1980) by *Bacillus subtilis* followed by (Nair and Pradeep 2002), who investigated lactic acid bacteria leading to the synthesis of 20-200 nm and are called nanocrystals. In the present study, a bacterial isolate *Bacillus licheniformis* strain GPI-2 was isolated from pebbles samples of a local gold mine near Khaltunala latitude (31.21515044oN), which synthesized gold nanoparticles of 20-35nm size and hexagonal as well spherical shape. This study was undertaken to evaluate the efficiency of gold nanoparticles synthesized by *Bacillus licheniformis* GPI-2 to control significant plant root nematode (*Meloidogyne incognita*).

*Meloidogyne incognita* is a nematode belonging to family *Heteroderidae* and is commonly known as root-knot nematode as it prefers to attack the root of its host plant. It has worldwide distribution and numerous hosts. When M. incognita attacks the roots of host plants, it sets up a feeding location, where it deforms the normal root cells and establishes giant cells. The roots become gnarled or nodulated, forming galls, hence the term “root-knot” nematode (Pline et al., 1988).

Tomato (*Solanum lycopersicum* L) is an important vegetable crop, and its cultivation occurs worldwide. Yield loss due to root-knot nematodes (*Meloidogyne* spp.) on tomato range from 40 to 46% in India (Perez et al., 2010). Management of *Meloidogyne incognita* is carried out by methods such as soil fumigation, nematicidal seed treatments, post-emergence nematicidal application, and cultivars partially resistant to *Meloidogyne incognita*. Tomato Plants infected with *Meloidogyne* spp. show typical symptoms of root galling. Keeping this in view present study was carried out to evaluate the nematicidal effect of gold nanoparticles on *Meloidogyne incognita*.

## Material and methods

### Isolation of bacterial isolate

The bacterial isolate used in this study was isolated from a gold mine of Khaltunala using serial dilution technique in nutrient agar after incubation at 37°C for 48 hrs. The bacteria were characterized morphologically, and biochemically and finally molecular using 16S rDNA technology (Holt et al., 2000). Qualitative determination of gold nanoparticles synthesizing ability of *Bacillus licheniformis* was determined which was depicted by the color change from pale yellow to red wine color.

### In vitro synthesis of gold nanoparticles by indigenous *Bacillus licheniformis* strain GPI-2

Extracellular biosynthesis of gold nanoparticles was carried out using supernatant of *Bacillus licheniformis* strain GPI-2, treated with 1mM gold chloride solution followed incubation at 37°C and to achieve maximum gold nanoparticles activity, the time range of 0-240 hrs was investigated.

### Characterization of gold nanoparticles was carried out using *Bacillus licheniformis* strain GPI-2

#### UV-vis spectroscopy

The biosynthesis of gold nanoparticles indicated via visual observation of the color change of the culture filtrate and confirmed by UV visible spectroscopy.

#### Fourier transform infrared (FTIR)

Microcup was washed with 100% absolute ethanol. 10 ul samples was filled in a 2 mm internal diameter micro-cup and loaded onto the FTIR set at 26°C ± 1°C. The samples were scanned in the range of 4,000 to 400 cm−1 using a Fourier transform infrared spectrometer (Thermo Nicolet Model 6700, Waltham, MA, USA). The spectral data obtained were compared with the reference chart to identify the functional groups present in the sample.

#### Transmission electron microscope

A drop of the sample was applied to a carbon coated copper grid. After about 1 min, the excess solution was removed using blotting paper and the grid was air dried before analysis.

### The direct vulnerability of Meloidogyne incognita

#### Germination of tomato seeds

To study the nematicidal effect the bio gold GNPs prepared as above *Solanum lycopersicum* were prepared from Departmental of Vegetable Science DR YSP UHF Nauni. Thus seeds were shown in pot tray containing autoclaved sand and topsoil (1:2) for germination. The seedlings after reaching two leaf stages were transplanted into plastic pots containing a similar mixture as above. In this manner, 60 seedlings were transplanted into 60 pots and these were maintained in the greenhouse to be used for further bioassay of nematode egg hatch and survival.

#### Meloidogyne incognita

The culture of *Meloidogyne incognita* was maintained on the tomato plants. They were maintained as a greenhouse stock culture on tomato cv. Rachna. Infected plants were uprooted, and the entire root system was dipped in water and washed gently to remove adhering soil. Egg masses of M. incognita were collected using forceps, disinfected with a 0.4% solution of sodium hypochlorite for three minutes then rinsed three times with sterilized water. Egg masses were placed in an autoclaved tissue grinder and gently crushed, then pipetted onto an autoclaved 25 μm aperture sieve and rinsed with sterilized water. A subset of the collected eggs was used for the egg hatch bioassay whilst the remainder was incubated at 25°C until hatched in order to obtain J2. The fresh nematode inoculum (2.00 J2ml-1) was used for in vitro and pot experiments.

The vulnerability of *M. incognita* to gold nanoparticles in water J2 of M. incognita isolated from tomato plants were used for this direct vulnerability assay to gold nanoparticles was done to get an insight of nematicidal activity of gold nanoparticles. 30 nematodes were added to 3ml of a solution containing 0,100,200,300,400,500 ul of a colloidal solution of GNPs with five replication treatment and incubated at room temperature 25OC. Nematode mortality rate was measured with an inverted microscope. After every 30 minutes, samples were checked for the mortality and their rate was recorded to determine the effective dose required for affecting nematodes. Healthy nematodes were defined as those were curled where as vulnerable or unhealthy nematodes were defined as those that appeared stiff or straight bodies.

#### Soil treatment with gold nanoparticles

The soil was inculcated with *M. incognita*. The water saturation level of the 50 cm3 soil sample was predetermined to be 25ml. The total was homogenized divided into 50 cm3, placed into a plastic container and then saturated with 25ml. gold nanoparticles solution at 0,300,600,900,1200,1500 ul. The samples were arranged in a completely randomized design with five replicates and incubated at room temperature for one to ten days. After the designated exposure time nematodes were extracted after from samples using Baermann tray system. After 48 hrs submergence in water, the samples were then poured into sieve filtered *M. incognita* were counted using an inverted compound microscope.

#### Pot plant experiment

Pot plant experiment with gold nanoparticles was carried out. Pots containing soil which was already infected with *M .incognita* culture and seedling of tomato were transplanted into pots and these pots were again inoculated with a fresh culture of *M. incognita* followed by application 0 to the 1500ul colloidal solution of GNPs to the pot plants. Plants wilting and others symptoms for diseases were checked and plant shoot lengths were recorded to the effect of GNPs on plants physiological processes and growth parameters. Treatments varied from C, T-1, T-2, T-3, T-4 and T-5 i.e 0,300, 600, 900, 1200,1500 µl of bio gold nanoparticles synthesized by *Bacillus licheniformis* strain GPI-2.

#### Estimation of photosynthetic pigments

Photosynthetic pigments (chlorophyll a) in leaves were assayed according to Hiscox and Israelstam (1979). The extraction was made from 100 mg of fresh sample in acetone (80%) in the dark at the room temperature and was measured with a UV/VIS spectrophotometer (Shimadzu UV-160, Kyoto, Japan) ( Salem and Amari, 2013)

#### Measurements of total leaf conductance and transpiration rate

Total leaf conductance and transpiration rate of the tomato measured using LiCor, 6400XT, Lincoln, NE, USA.

## Results

Isolation of gold nanoparticles synthesizing bacteria was carried out from different samples viz. yellow soil, pebbles, biofilm and stalagmites collected from a local gold mine using nutrient agar medium at 37ºC (Figure-1). Which was identified as *Bacillus lichniformis* GPI-2 after morphological, biochemical and molecular characterization using 16 S rDNA technology.

**Figure 1:**
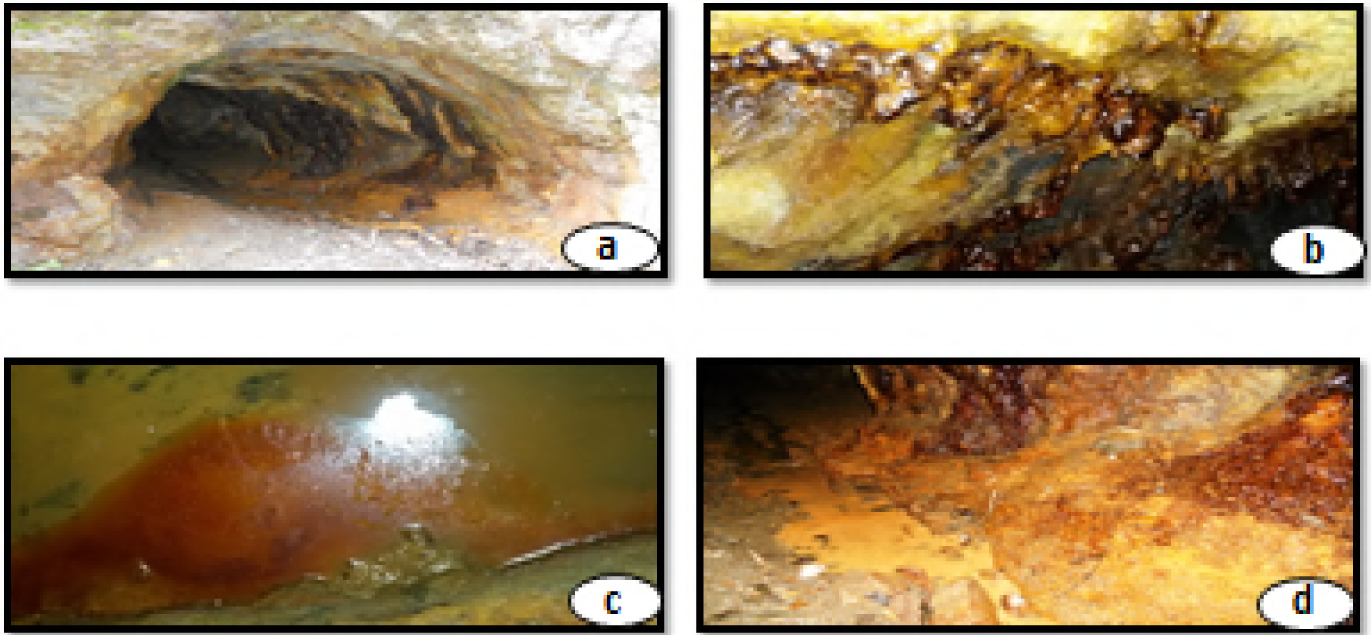
Different sites of samples collection of gold mine Khaltunala (a) pebble (b) roof topping (c) biofilm (d) yellow soil

### In vitro synthesis of gold nanoparticles by indigenous *Bacillus lichniformis* strain GPI-2

Extracellular biosynthesis of gold nanoparticles was carried out using a supernatant of *Bacillus licheniformis* strain GPI-2, treated with 1mM gold chloride solution and incubated at 37°C for a time period of 0-240hrs. The biosynthesis absorption spectra of gold nanoparticles which were indicated by the color change of solution from yellow to red wine (Figure-2) and was further confirmed by spectrophotometrically. The gold nanoparticles synthesis activity increased with the increase in incubation period up to 36 hrs and removed stable up to 48 hrs (Figure-3) and the time of incubation course and increase information of gold nanoparticles took place up to 36 hrs and remained stable up to 48 hrs and then the values declined up to 240 hrs.

**Figure: 2.**
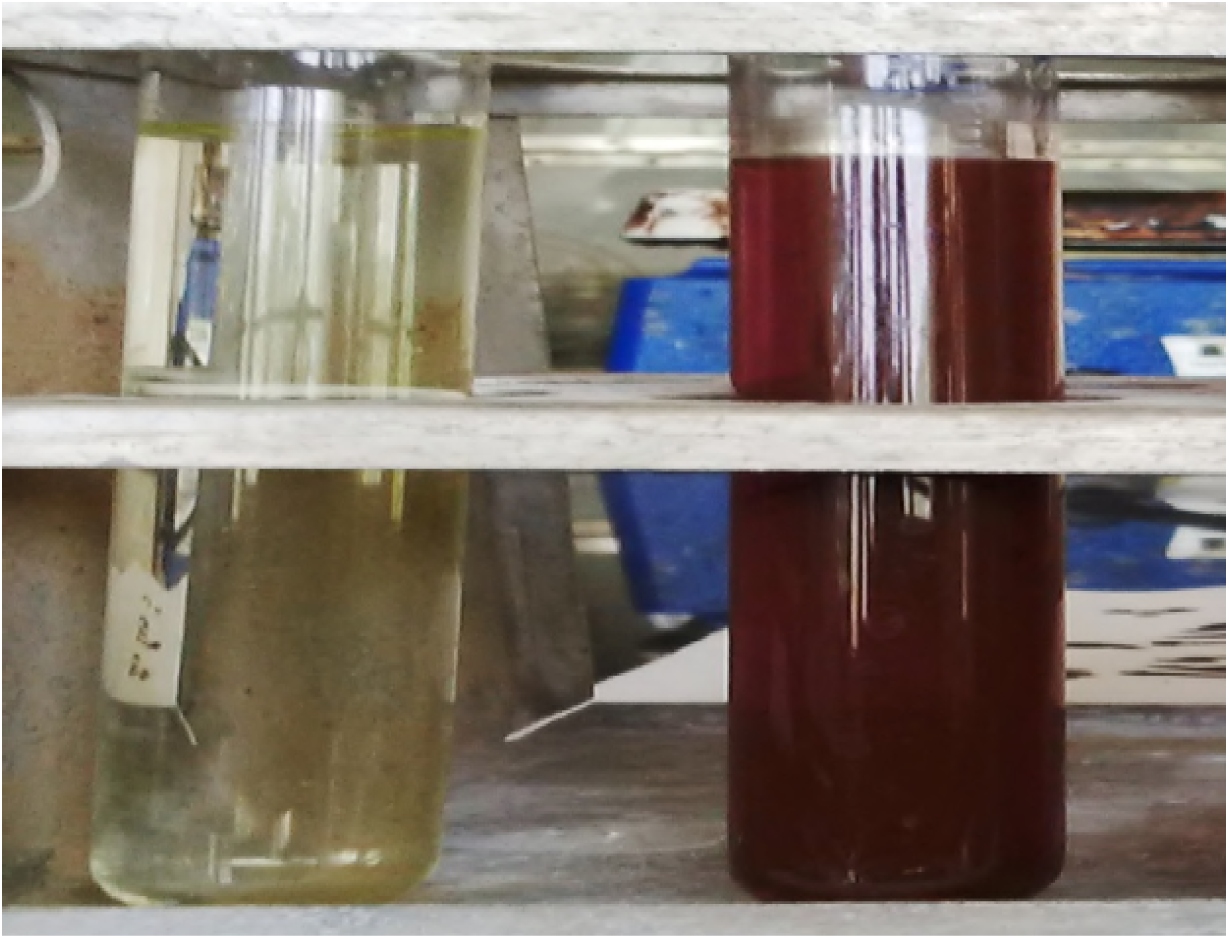
Biosynthesis of gold nanoparticles by *Bacillus lichniformis* GPI-2

**Figure: 3.**
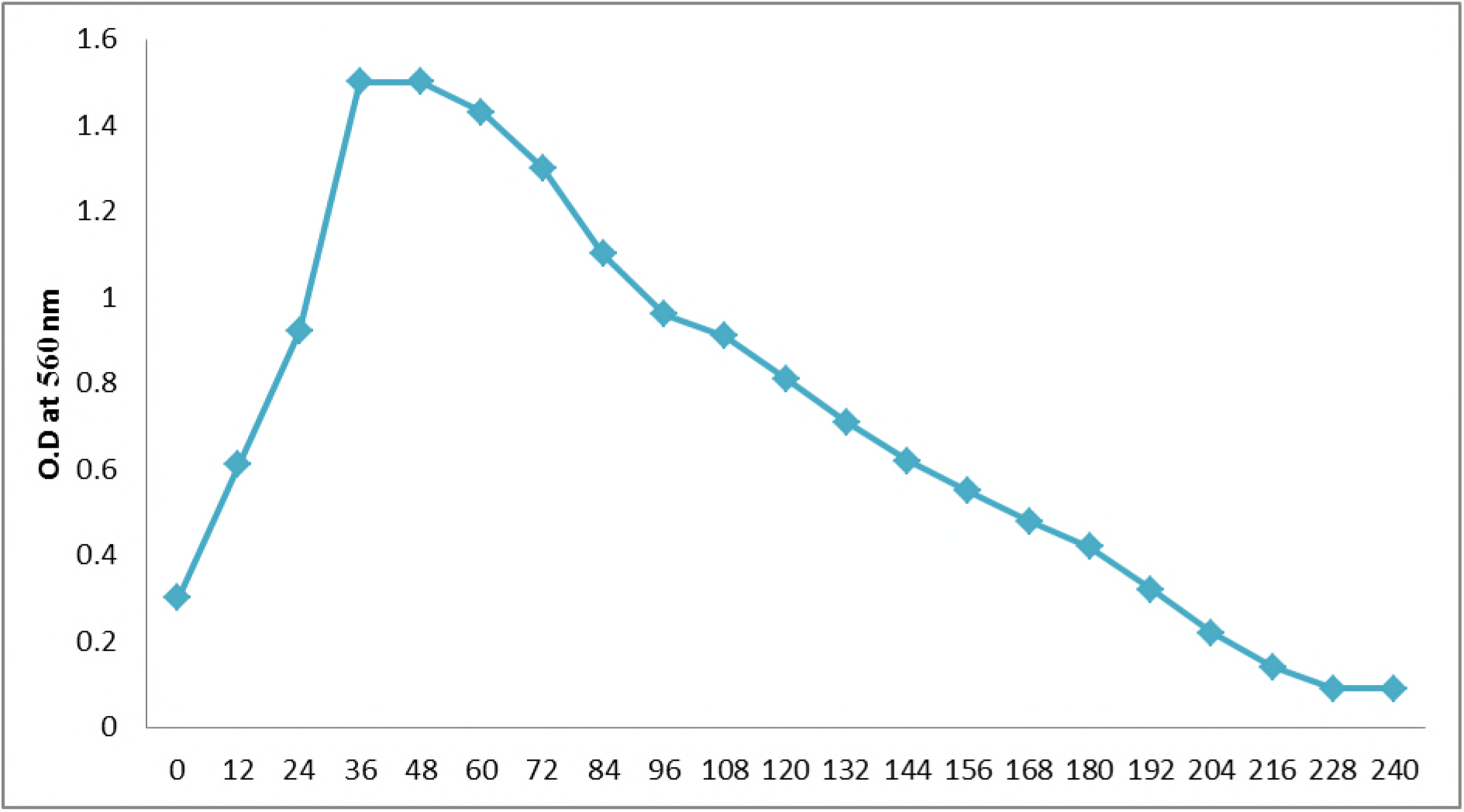
UV-Vis absorption spectra of invitro synthesised gold nanoparticles after incubation of *Bacillus lichniformis* GPI-2 with 1mM gold chloride for a time period of 0-240 hrs at pH: 6.8

### Characterization of in vitro synthesis of nanoparticles by *Bacillus lichniformis* strain GPI-2

The TEM image (Figure-4) clearly showed discrete gold nanoparticles in the size range of 20 to 45 nm which were mostly triangular, irregular and hexagonal indicating that it was possible to synthesize gold particles of nano dimensions with a satisfactory level of monodispersity. FTIR measurements were carried out to identify the possible biomolecules protein responsible for the capping and efficient stabilization of the gold nanoparticles synthesized by *Bacillus lichiniformis* GPI-2. FTIR spectrogram has shown the presence of four peaks 3280.18, 2380.99, 2109.12 and 1636.32 (Figure-5). High sensitivity to small variations in molecular geometry and hydrogen bonding patterns makes the amide I band uniquely useful for the analysis of protein secondary structural composition and conformational changes (Wright, 1947). The FTIR spectra reveal the presence of different functional groups. Wave number between 3235 and 3280 cm-1 indicates for hydrogen bond lengths between 2.69 to 2.85Ao. Alkynes C-C triple bond stretch is found at 2109 cm-1. A peek at 1636 cm-1 corresponds to the N-H bend of primary amines due to carbonyl stretch. Amide 1 is most intensive absorption band in protein. It is primarily governed by the stretching vibration of the C=O (70-80%) and CN stretching groups. (10-20%) frequency 1600-1700 cm-1. In the amide, I region (1700−1600 cm-1), each type of secondary structure gives rise to a somewhat different C=O stretching frequency due to unique molecular geometry and hydrogen bonding pattern. N-H Stretch of primary and secondary amines, amides. Amide A is with more than 95% due to N-H stretching vibration. This mode of vibration does not depend on the backbone conformation but is very sensitive to the strength of a hydrogen bond. Peak maximum around 1650 cm-1 corresponds to proteins alpha-helical structure. The half width of the alpha helix band depends upon on the stability of the helix. When half width of about 15 cm-1 then we have more stability of helix and transition free energy of more than 300 cal/ml. Amide 1absorption is primarily determined by the backbone conformation and independent of the amino acid region, its hydrophilic or hydrophobic properties and charge. The average frequency of the main compounds is about 1629 cm-1. Arginine amino acid role was found at 1636 cm-1 in gold nanoparticles synthesis through results of FTIR.

**Figure 4:**
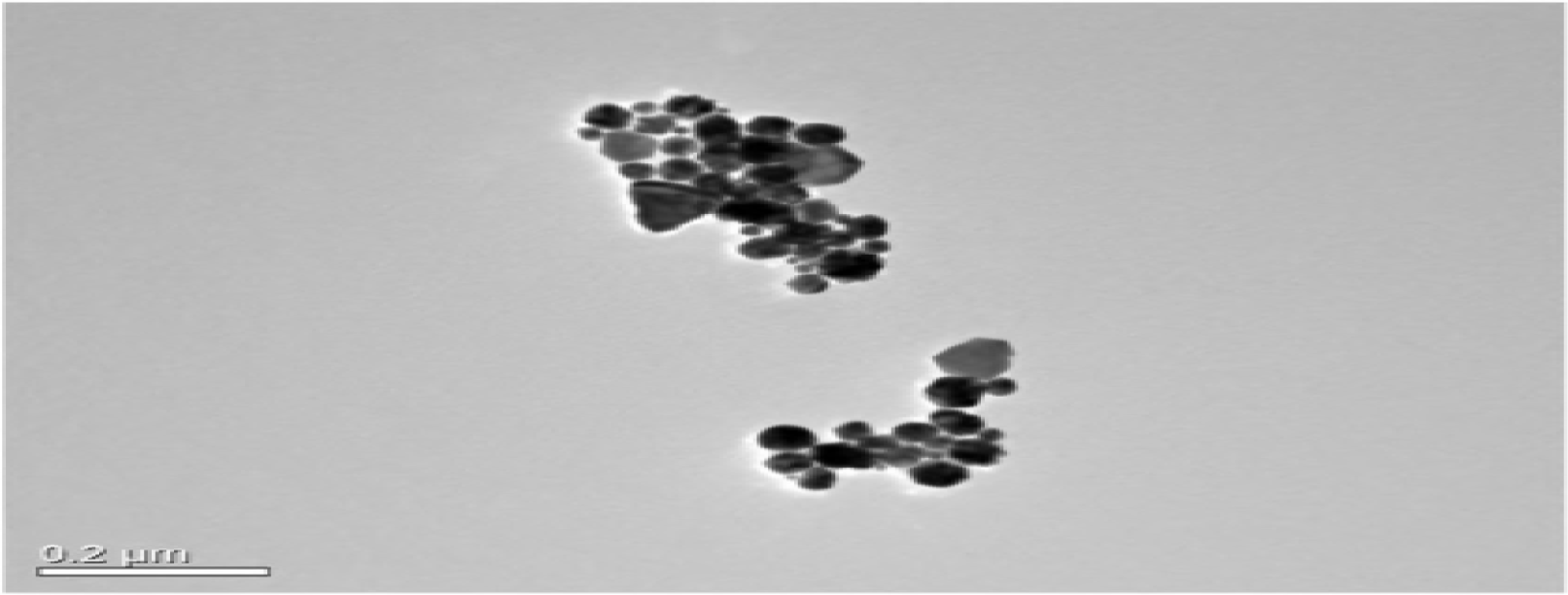
Characterization of gold nanoparticles through transmission electron microscope showing the different morphology of gold nanoparticles.

**Figure: 4.**
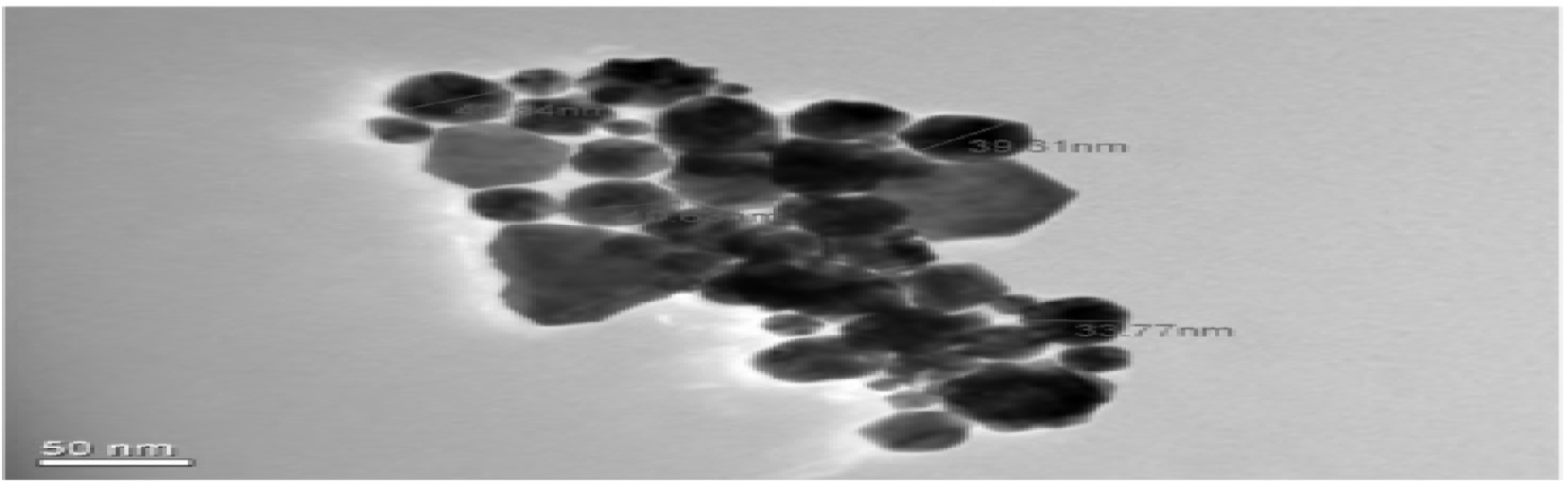
TEM image of gold nanoparticles showing different sizes of gold nanoparticles

### The direct vulnerability of *Meloidogyne incognita* to gold nanoparticles in cavity block experiment

The direct vulnerability of *Meloidogyne incognita* to gold nanoparticles in water was evaluated and it was found that gold nanoparticles increased the mortality of *Meloidogyne incognita* effectively and efficiently. After one hour of application of GNPs mortality was initiated and 100% mortality has been achieved after three hours. It was observed that after application of gold nanoparticles crystallization of water occurred even at room temperature followed by clumping or aggregation of these nematodes leading to mortality. T-4 treatment consists of 400µl of bio gold nanoparticles found to be highly effective to cause the maximum mortality whereas in case of control no mortality has been found (figure-5a, b).

**Figure 5a.**
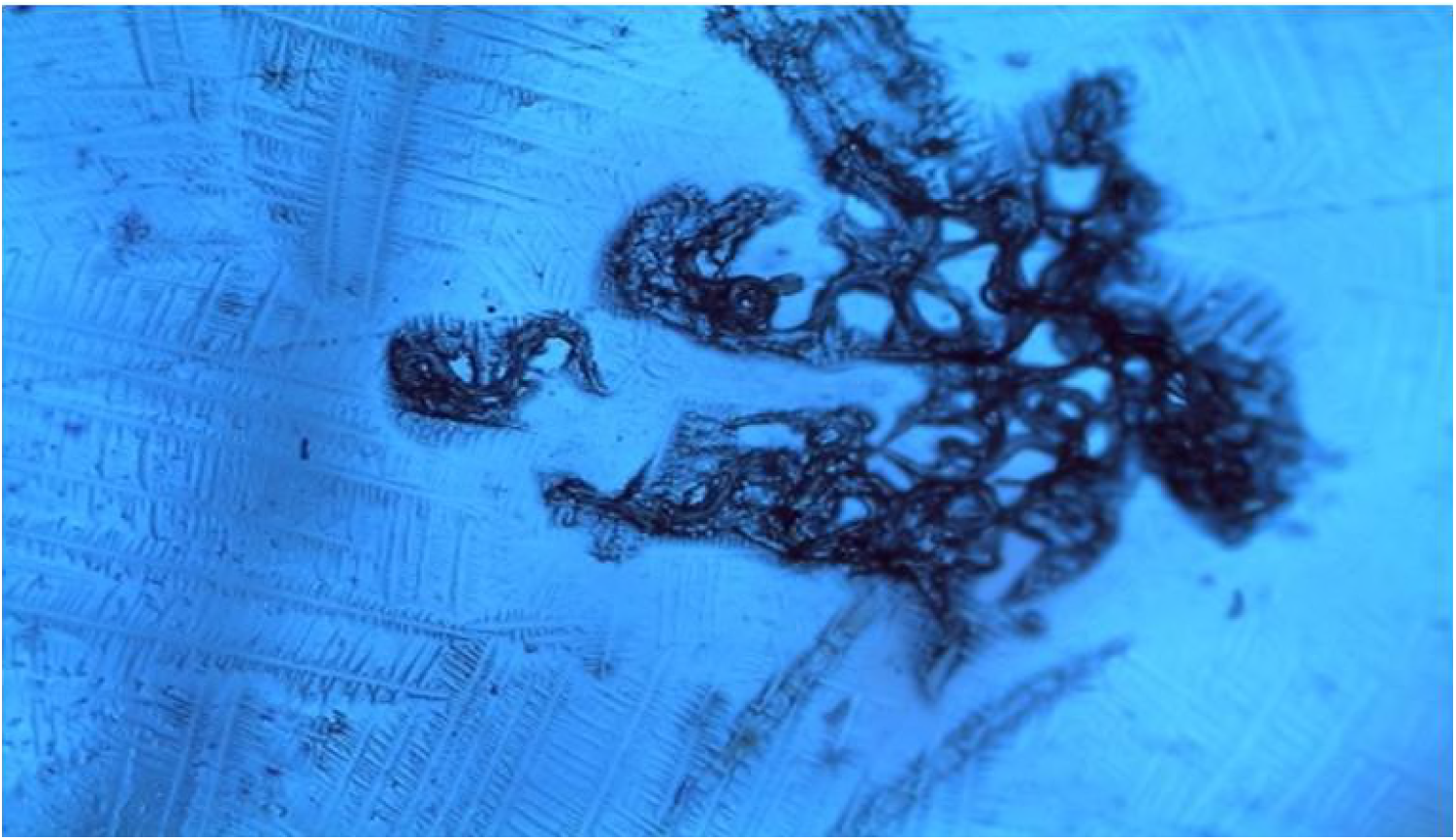
*M. incognita* root knot nematodes were found to be killed after applying 100-500ul of gold nanoparticles. 400 ul dose was found to be highly effective. where 100% lethality was found in just three hours. It has been observed that gold nanoparticles causes crystallization of water and have adverse effect on survivability of *M. incognita*.

**Figure -5b.**
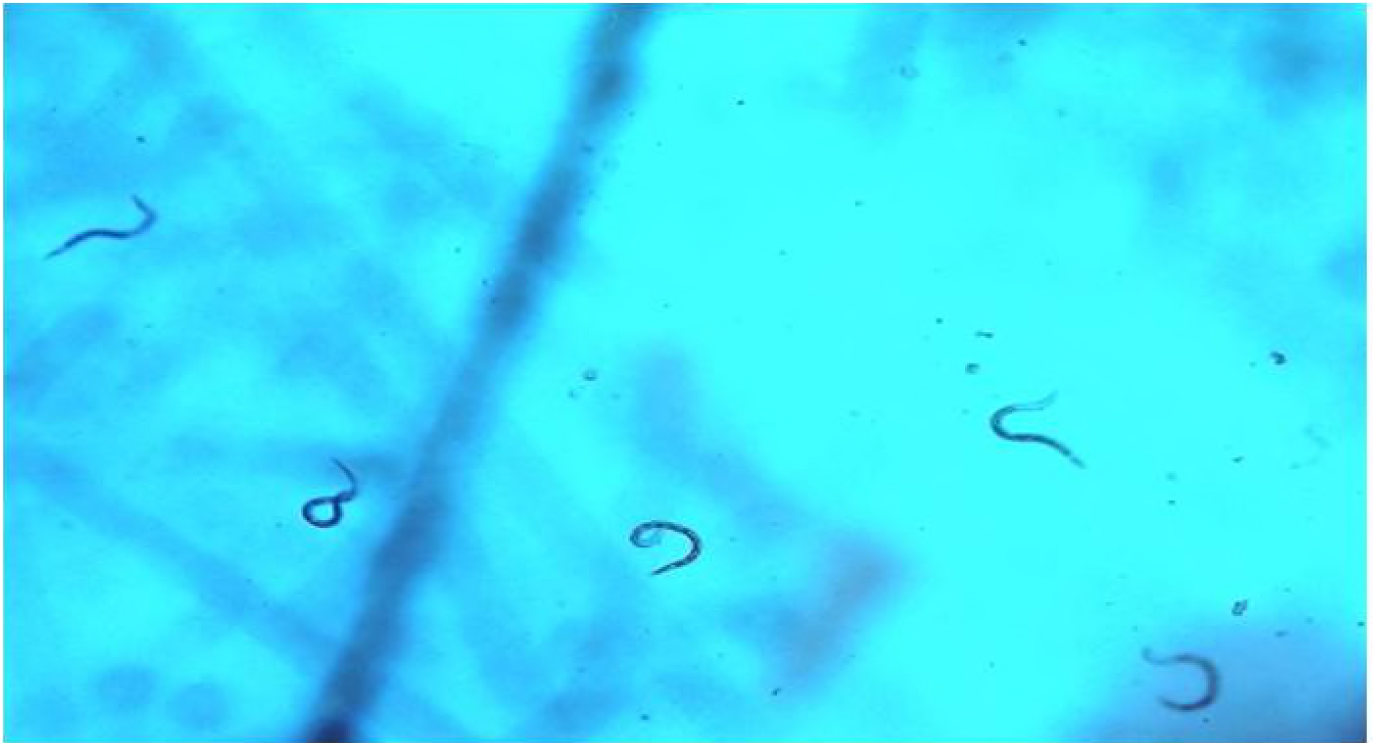
*M. incognita* healthy culture before applying gold nanoparticles.

### Soil treatment with gold nanoparticles

#### Soil treatment with gold nanoparticles

Enumeration of *M. incognita* extracted from soil samples was found to be reduced significantly treated with gold nanoparticles in a repeated laboratory experiment. The exposure time of 6-8 hours did not affect the number of *M. incognita* and during this time there was no significant effect of GNPs on *M. incognita* population in the soil. There was an interaction between exposure time and GNPs. Initially, there was a linear relation of time with a number of unhealthy nematodes. As exposure time increased death of nematodes increased drastically. The number of *Meloidogyne incognita* extracted from soil samples were reduced when the soil samples were treated with GNPs with different concentration.0, 100, 200, 300, 400ul GNPs. Most effective treatments were 300 ul and 400 ul because they have maximum mortality rate (Figure-6). GNPs reduced the number of J2 recovered after 1to 4th day of exposure of gold nanoparticles.

**Figure 6:**
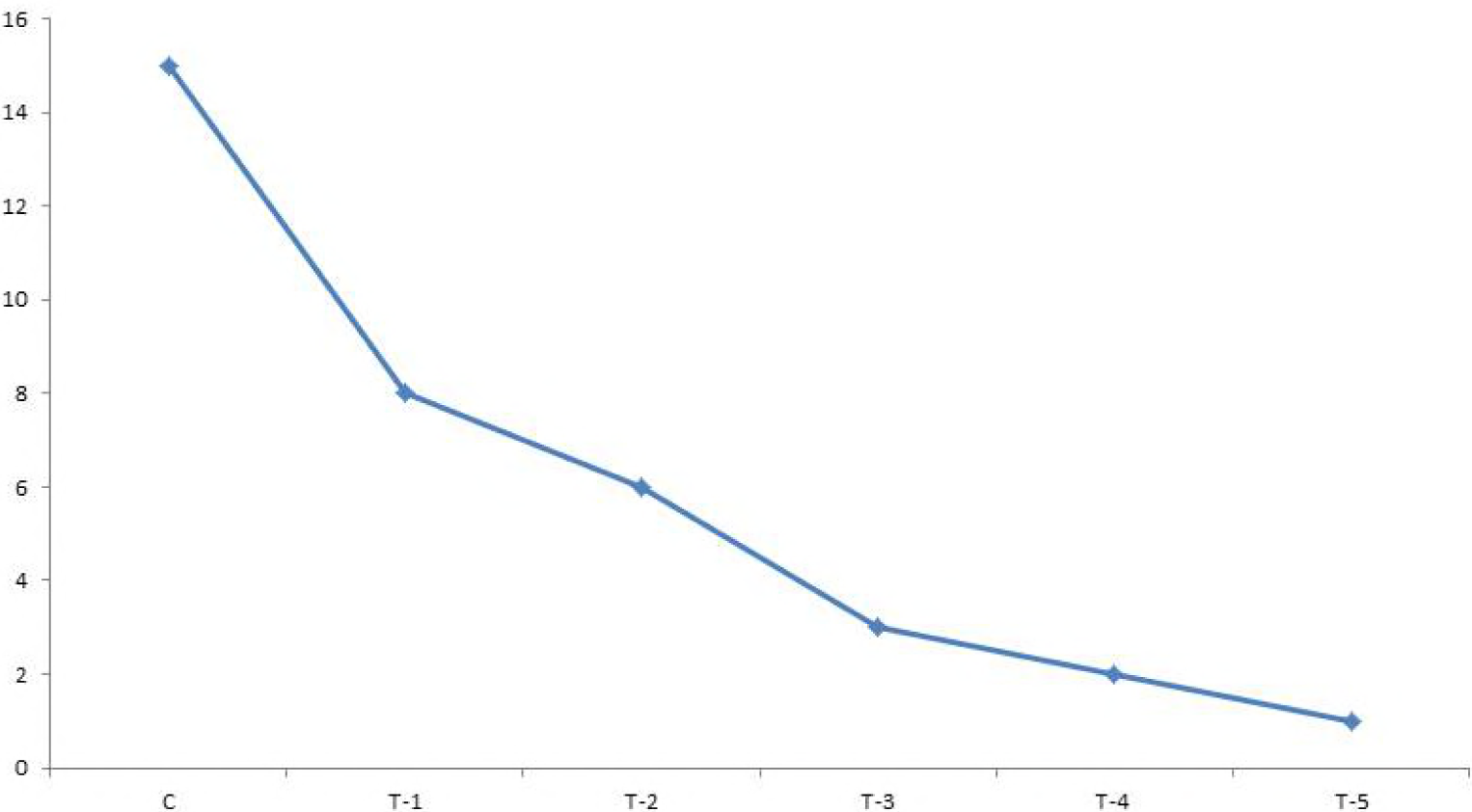
Number of *Meloidogyne incognita* extracted from soil samples were reduced when the soil samples were treated with GNPs with different concentration. 0, 300,600,900,1200,1500 ul GNPs. Most effective treatments were 1200 ul and 1500ul because they have maximum mortality rate.

#### Evaluation of different concentration of the bio gold nanoparticles on *M.incognita* infected tomato seedlings.

Tomato seedlings were raised from seeds purchased from Dr. Y.S Parmar University of Horticulture and Forestry Nauni. Tomato seedlings were infected with the live culture of *M. incognita* followed by application of different concentration of biogold in case of control no GNPs were applied but in T-1, T-2, T-3, and T-4 GNPs were applied. Which ranges from 0 to 400ul of GNPs. Higher doges of GNPs were highly effective to prevent the infection *M. incognita* to tomato plants; it also found that gold nanoparticles have increased the shoot growth of tomato plants (Figure-7). GNPs application in pot plants has shown higher resistance to M. incognita infection whereas control plants found highly susceptible to *M. incognita* infection.

**Figure 7:**
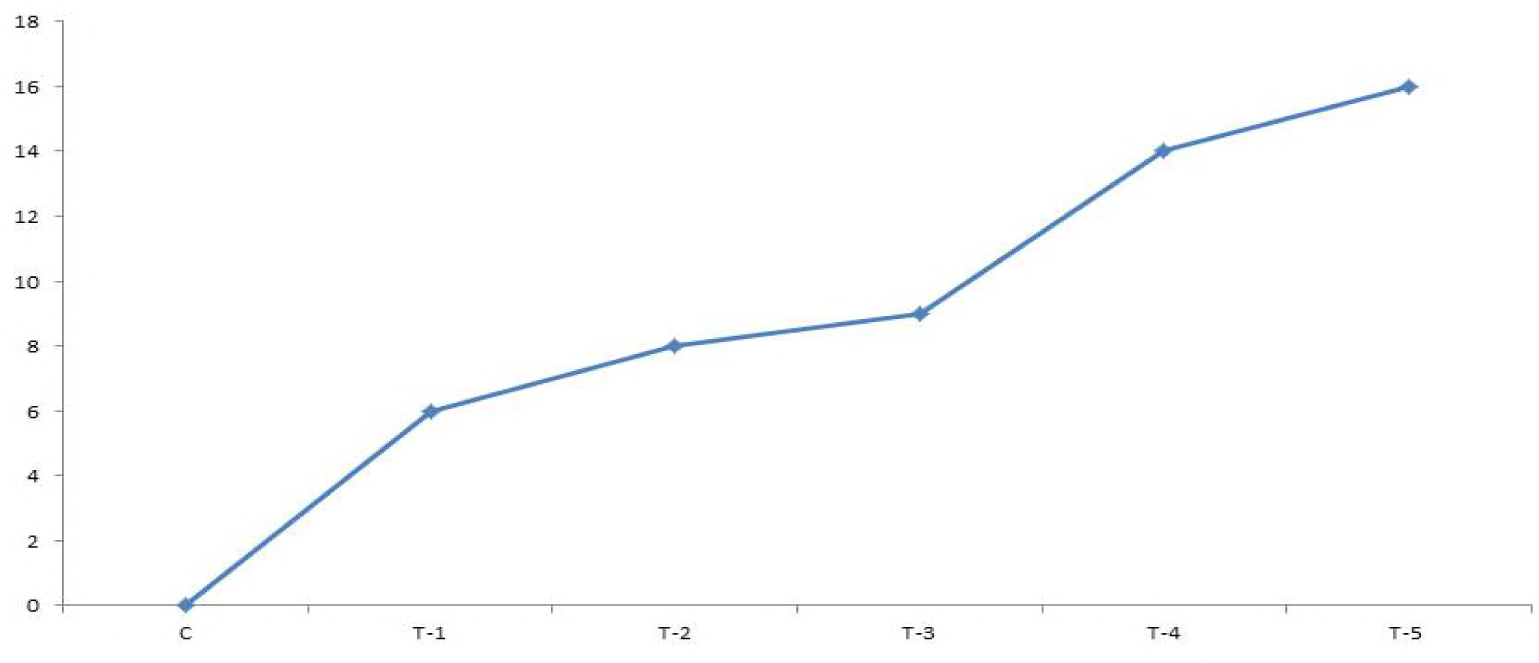
*M .incognita* mortality caused after foliar spray of gold nanoparticle application. It has been found that higher doges of GNPs were highly effective to prevent the infection *M .incognita* to tomato plants at T_4_, T_5_

### Plant growth promoting

#### Effect of gold nanoparticles of *Bacillus sonorensis* KRI-7 on seed germination and plant growth

Both in-vitro and in vivo experiments were conducted to study the effect of gold nanoparticles synthesizing by *Bacillus sonorensis* KRI-7, on seed germination and plant growth of *Spinacia oleracea* and *Solanum lycopersicum*. Seed germination under in-vitro conditions has been found to occur earlier in gold nanoparticles treated seeds as compared to untreated seeds by one day. It has also been found that in *Spinacia oleracea* percent seed germination increased significantly with T-1 (80%) treatment (100µl) of bio gold nanoparticles of *Bacillus sonosensis* KRI-7 as compared to control (56%). However, percent germination decreased to 64% when treated with 200µl of bio gold nanoparticles of *Bacillus sonosensis*. In case *Solanum lycopersicum* seed germination increased with increase in the concentration of bio gold nanoparticles of *Bacillus sonorensis* KRI-7. It has been observed that only 36% seed germinated in control samples. Seeds treated with 100 µl of biogold nanoparticles showed 58% germination and seed germination increased to 66% in T-2 treatment. The result has been obtained with respect of tomato with increased fresh weight by T-2 treatment following by T-1 treatment as compared to control. The dry weight data was also found to be similar with fresh weight data (Figure-20a, b). The results obtained from the in vivo experiment effect of biogold nanoparticles *Bacillus sonorensis* KRI-7 of both plant species was found to produce a very significant increase. In case of *Solanum lycopersicum* shoot lengths increased by one cm in T-1 treatment as compared to control value of 6 cm and with the T-2 treatment of bio gold nanoparticles, shoot length increased to 9 cm which is a significant increase. The similar result has been obtained an increase in leaf area of tomato. The control samples leaf area showed 4 cm^2^ which increase to 5 cm^2^ with T-1 treatment and 6 cm^2^ with T-2 treatment (Figure-7b).

**Figure 7b:**
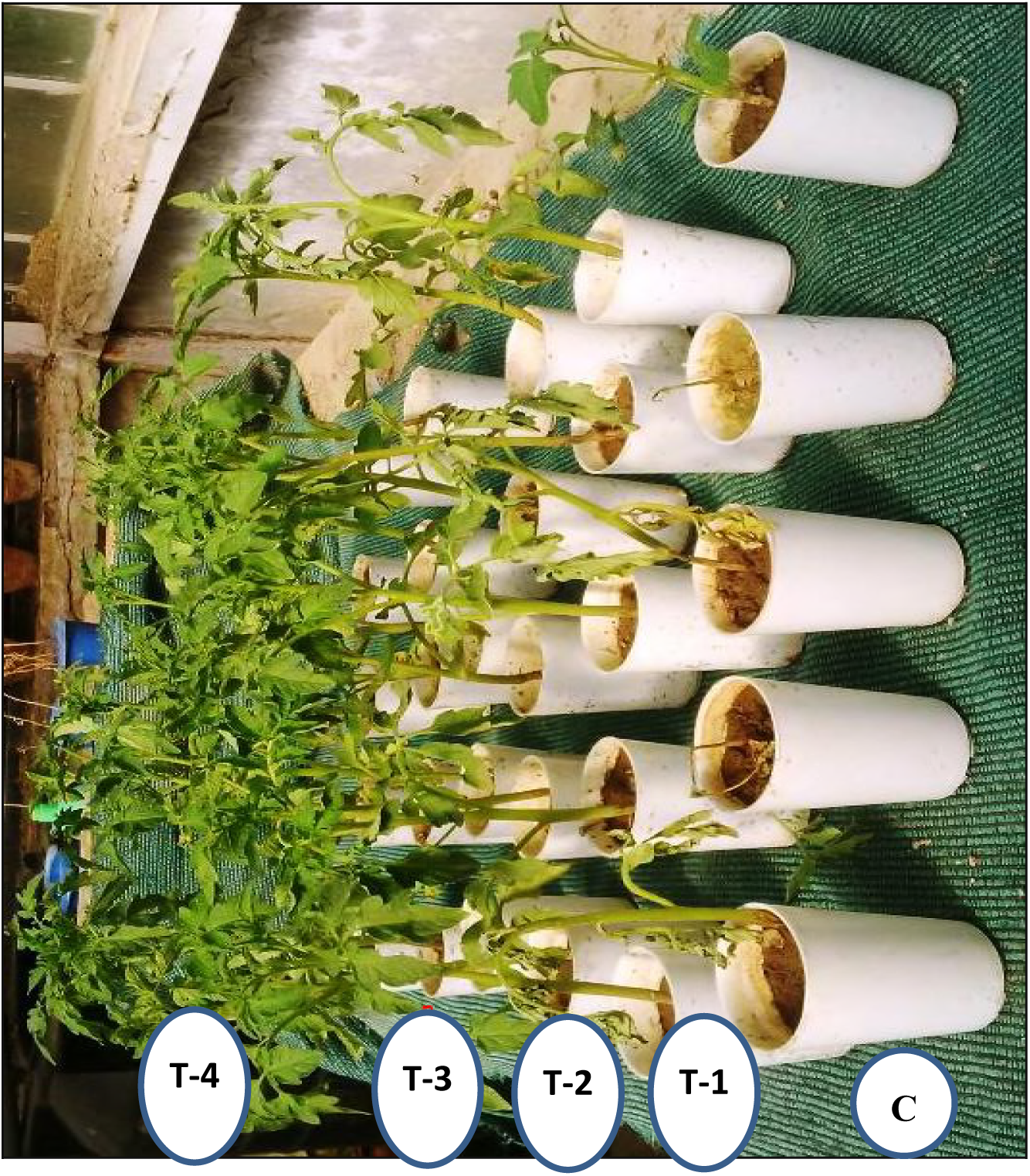
*M. incognita* treated tomato seedlings C: without gold nanoparticles T1: Treatment with 100µl of biogold nanoparticles T2: Treatment with 200µl of biogold nanoparticles T3: Treatment with 300µl of biogold nanoparticles T4: Treatment with 400µl of biogold nanoparticles

#### FTIR analysis of Solanum lycopersicum

Fourier Transform Infrared Spectrophotometer (FTIR) is perhaps the most powerful tool for identifying the types of chemical bonds (functional groups) present in compounds. The wavelength of light absorbed is characteristic of the chemical bond as can be seen in the annotated spectrum. In case of *Solanum lycopersicum* peaks were founding at which 3860, 3536, 3283, 3327, 2921, 2854, 2169, 2346, 1156, 1086 correspond to low concentration alcohols and phenols, low concentration of carboxylic acid, strong primary amines and conjugated nitrites, aromatic ether. These peaks have also been found in T-2 treated plants leaf, which indicated that gold nanoparticles did not have any negative impact on functional groups in case *Solanum lycopersicum*. From these results, it is concluded that gold nanoparticles did not affect functional groups in *Solanum lycopersicum* and thus these are safe to be consumed (Figure-7c).

**Figure 7c:**
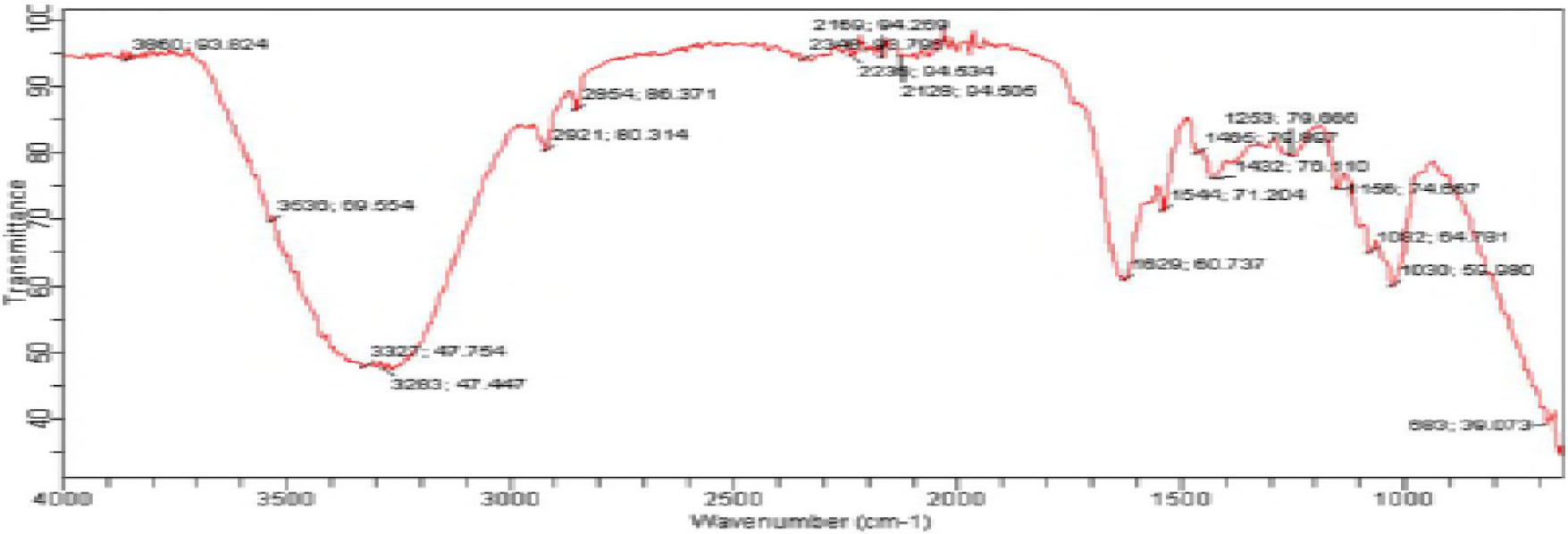
FTIR spectrum of gold nanoparticles generated by *Bacillus licheniformis* strain GPI-2

In the present investigation, we have found that gold nanoparticles significantly increased the seed germination and shoot length of tomato seedlings in comparison to control plant 21.67 percent increase in the shoot length have been found in case of T-2 treated plants. Jha et al. (2009) reported the influence of gold nanoparticle fenugreek (*Trigonella foenumgraceum* L) seedlings and assessed the effect of gold nanoparticles on leaf number, root length, shoot length and net weight and diosgenin content. Deka et al. (2015) studied the influence of gold nanoparticles lentils seeds and found that seed germination index, root length, shoot length and other parameters increased by gold nanoparticles. It was also that accumulation and uptake of nanoparticles is dependent on the exposure concentration. Jha et al. (2009) investigated the effect of gold, copper and iron nanoparticles on germination and seedling vigor index of wheat seeds and concluded that seed treatments and incubation time affected seedling growth such as root and shoot growth. Seedling vigor increased when seeds were soaked in nanoparticles. It was reported toxicity of gold nanoparticles on *Brassica Juncea* var.varuna. Gold nanoparticles induced a slight increase in the root, shoot length, chlorophyll content and protein content. Deplanche et al. (2007) examined the impact of gold nanoparticles on the physiology and nutritional quality of radish sprout and suggested that gold nanoparticles could significantly affect the growth and nutrient content, confirmation in radish sprouts.

#### Photosynthesis, transpiration, conductance

In our study, we have observed that gold nanoparticles treated tomato plants have higher chlorophyll content as compared to untreated control plants and our other experiments like conductance and transpiration also have a similar effect on tomato plants. In the case of photosynthesis T4 treatment of GNPs provided the best result among all treated and untreated plants. (Figure 8, 9, 10)

**Figure 8:**
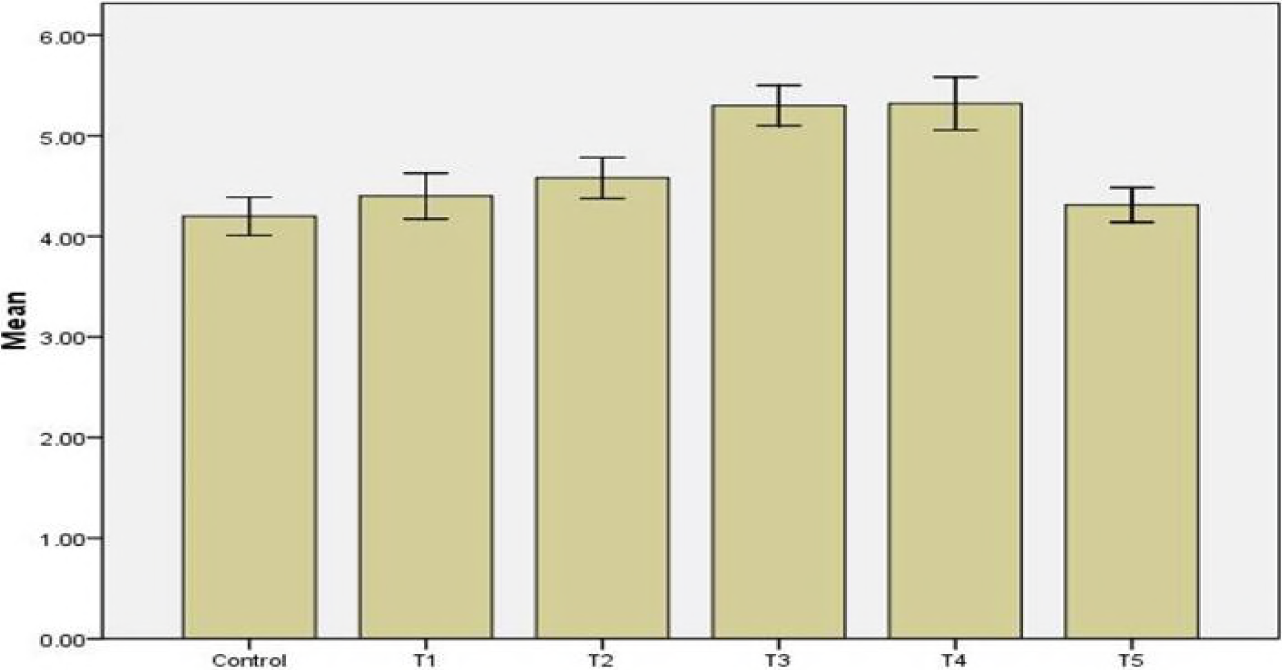
Gold nanoparticles mediated growth profile of tomato plants up to one week. Gold nanoparticles caused an increase in plants height, as compared to the untreated seedlings.

**Figure 9:**
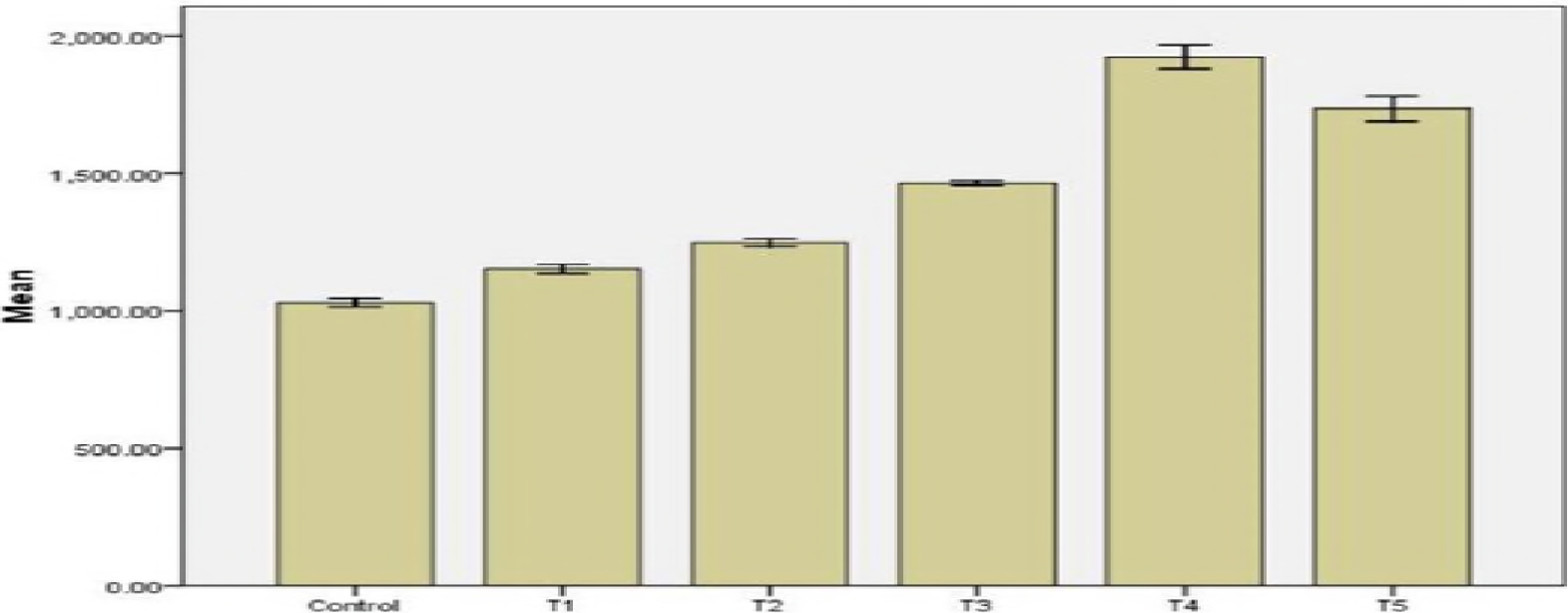
Effect of gold nanoparticles on chlorophyll component on tomato plants, it has been identified that T_4_ treated plants with 4ml of GNPs showed higher chlorophyll content than others.

**Figure 10:**
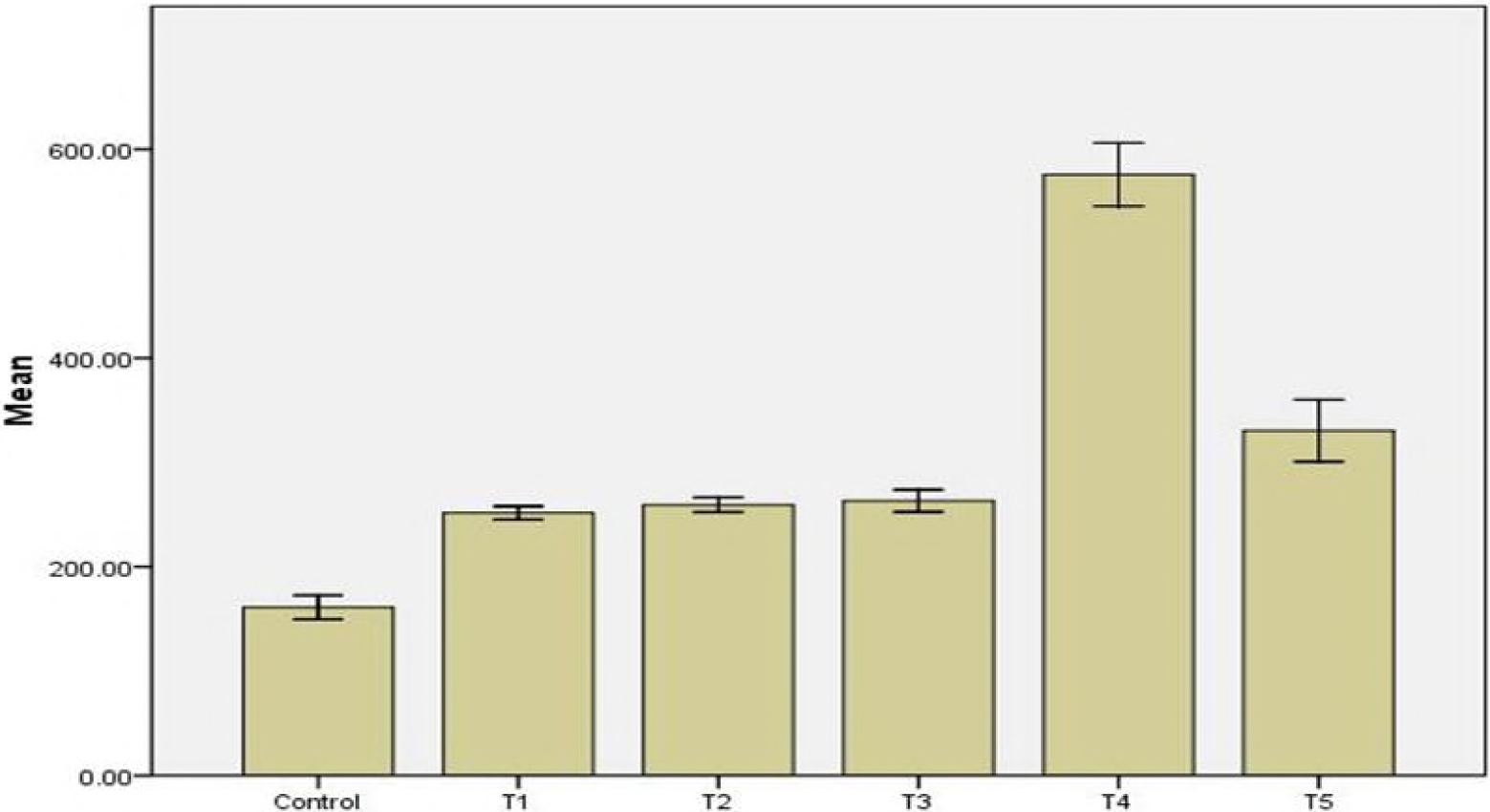
Effect of GNPs on transpiration rate, transpiration rate found to be higher in case of T_4_ plants.

**Figure 11.**
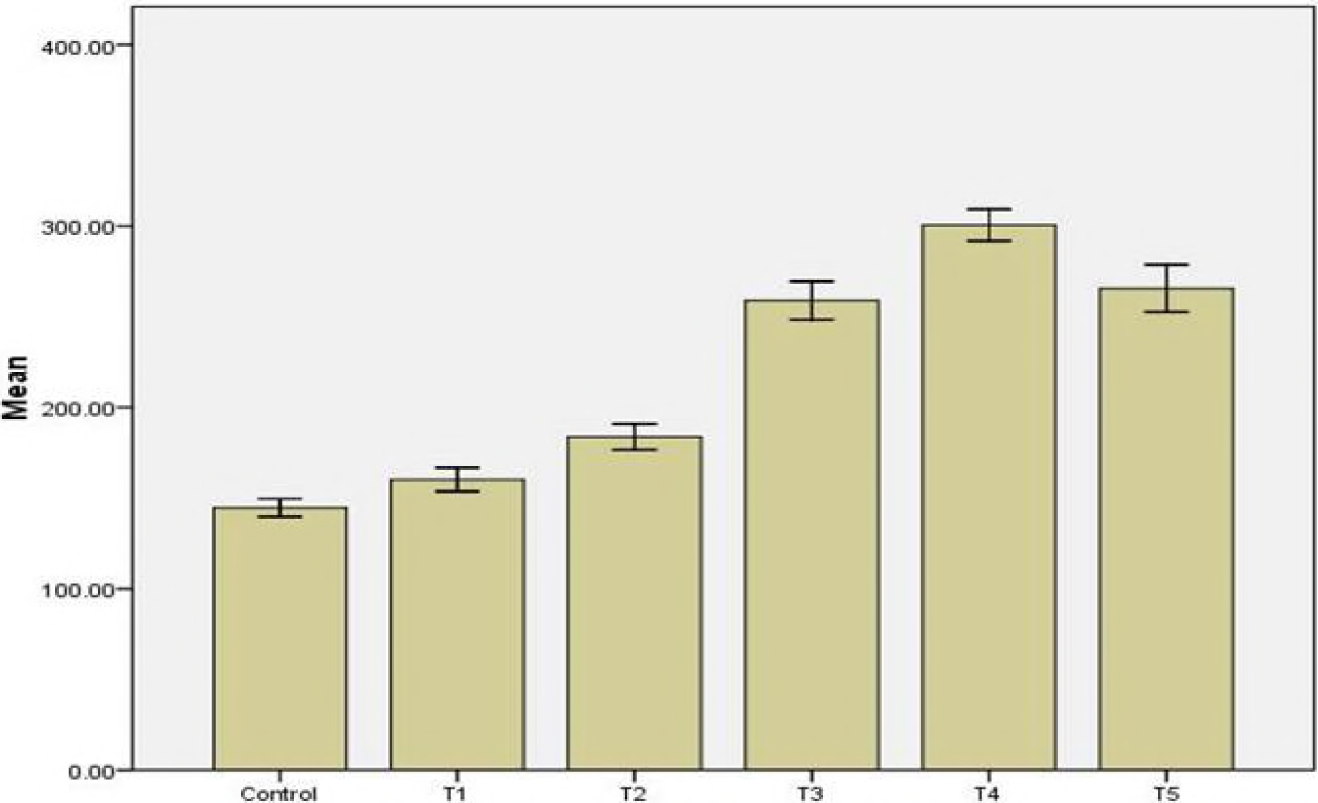
Effect of gold nanoparticles on conductance rate of tomato plants. T_4_ plants have shown the higher conductance than other treated as well control plants.

#### Effect of GNPs on nutrient uptake

Numerous studies have shown that uptake of nutrient elements is affected by the size of nanoparticles and the nature of species. Smaller the size, larger the surface area so their capacity to take more payload was increased, movement of nutrients (macro elements and microelements) foster by these nanoparticles. Where they move inside roots they carry a payload of different macroelements and microelements which significantly increase the growth of roots, shoots and make the better availability of nutrient of plants then untreated control plant.

## Discussion

Among the ambiance of natural resources, prokaryotic bacteria have received attention for the synthesis of gold nanoparticles. The reason for the bacterial preference for gold nanoparticles synthesis is the relative ease of manipulation. Microbes in general and bacteria, in particular, are preferred because of ease of downstream processing (Kashefi et al., 2001). Both gold thiosulfate and gold chloride have been reportedly used for the accumulation of gold by bacteria. The bacteria get killed, resulting in the release of organic molecules causing further precipitation of gold (Southam and Beveridge, 1996). The reduction and precipitation of gold involve periplasmic hydrogenases or cytoplasmic hydrogenases. Cytochrome C3 could be the complementary mechanism of gold reduction in bacteria. In the present investigation, gold chloride (HAuCl4), which is very hygroscopic and highly soluble in water, has been used successfully for the formation of gold nanoparticles. In the present study, for bioprospecting of gold nanoparticles producing bacteria-local gold mine and four hot water springs of Himachal Pradesh were investigated. Nangia et al., (2009) also described the isolation of gold nanoparticles synthesizing bacteria *Stenotrophomonas maltophilia* from singhbhum gold mines (Jharkhand). Birader and Lingappa (2012) were also successful for isolation of gold nanoparticles synthesizing *Bacillus* sp. from Hatti region of Karnataka. Some strains of *Arthrobacter* genera with the ability to synthesize gold nanoparticles have been isolated from basalt rock from Georgia (Kalabegishvili et al., 2012), Correa-Llanten et al., (2013) have stated the isolation of gold nanoparticles synthesizing *Geobacillus* sp. from Deception Island, Antarctica.

The present investigation has provided evidence that gold nanoparticles have the highest rate of mortality and were effective for management of root-knot nematodes (Cromwell et al., 2014) reported J2 of M. incognita were exposed to AgNPs in water at 30 to150ul/ml, 99% nematodes became inactive in 6 hrs. (Taha et al., 2016) evaluated the silver nanoparticles effect on EPN dependent on nano-Ag, concentration and exposure time. They have found significantly effect on EPNs reproductivity at two concentration 500 and 1000 ppm. Abdellatif et al., 2016 reported effective control of root Knot nematodes T. turbinate and *U. lactuca* similar to chemical control in eggplant.

Taha evaluated the nematicidal effect of silver nanoparticles on J2 *M. incognita* in the laboratory and in the screen house. When the *M. incognita* population was exposed to AgNP in water at 20, 40, 200, 500 and 1500ppm/ml achieved 96.5 percent mortality after 72 hrs with 1500 ppm. The concentration of 200 ppm caused 52 percent mortality on the third day and 1500 ppm was found to be the most effective dose while other doses were ineffective. Ardakani et al., 2016 studied effectiveness of AgNPs on M. incognita in case of tomato plants, dose of 800,400 and 200 mg/ml-1 of AgNP were highly significant to immobility and to mortality of J2 M. incognita and Ti02 nanoparticles showed 4.3 percent and 2 percent mortality when applied 800 and 400 mg/ ml-1 concentration. Abdellatif et al., 2016 studied the influence of silver nanoparticles on *M. javanica* reproduction and growth. They have observed that AgNPs treatment was equally effective as chemical treatment (Vydate 24%L), resulted in a reduction of egg-masses number per root system. The concentration of 17 mg, 100 mL-1 of U.lactuca with silver nanoparticles most effective in reduction of *M. javanica* population (69.44 percent J2s in the soil, number of females of *M. javanica* in roots reduced to 84.51 percent when treated with silver nanoparticles.

### Plant growth promoting

In the present investigation we have found that gold nanoparticles significantly increased the seed germination and increased shoot length of tomato seedlings in comparison to control plant 21.67 percent increase in the shoot length have been found in case of T4 treated plants and it has been found that smaller sized nanoparticles were more efficient for inducing growth. Jasim et al., 2016 investigated the influence of silver nanoparticle fenugreek (*Trigonella foenumgraceum* L) seedlings. They have assessed the effect of silver nanoparticles leaf number, root length, shoot length and met weight as well on the diosgenin content. Hojjat et al., 2016 studied the influence of silver nanoparticles lentils seeds. They have found that seed germination index, root length, shoot length, and other parameters were affected by silver nanoparticles. They have observed that accumulation and uptake of nanoparticles as dependent on the exposure concentration.

Yasmeen et al., 2015 investigated the effect of silver, copper and iron nanoparticles on germination and seedling vigor index of wheat seeds. They have concluded that seeds treatment and incubation time affect the seedling growth such as root and shoot growth. Seedling vigor increased when seeds were soaked in nanoparticles and incubated in distilled water. Pandey et al., 2014 studied toxicity of silver nanoparticles on Brassica Juncea var.varuna. Silver nanoparticles induce a slight increase in the root, shoot length, chlorophyll content and protein content. In our study, we have observed that gold nanoparticles treated tomato plants have higher chlorophyll content as compared to untreated control plants and our other experiments like conductance and transpiration also have a similar effect on tomato plants. In the case of photosynthesis T4 treatment of GNPs provided the best result among all treated and untreated plants. Nubia et al., 2016 examined the impact of silver nanoparticles on the physiology and nutritional quality of radish sprout and suggested that nAg could significantly affect the growth, nutrient content and macromolecules conformation in radish sprouts. When the seedling was exposed to 500 mg\L had 901mg AG\Kg dry weight and significantly less ca, Mg, B, Cu, and Zn, compared with control. They also revealed that changes occur in lipids, proteins and structural components of plants cell such as lignin, pectin, and cellulose.

**Table: 1.**
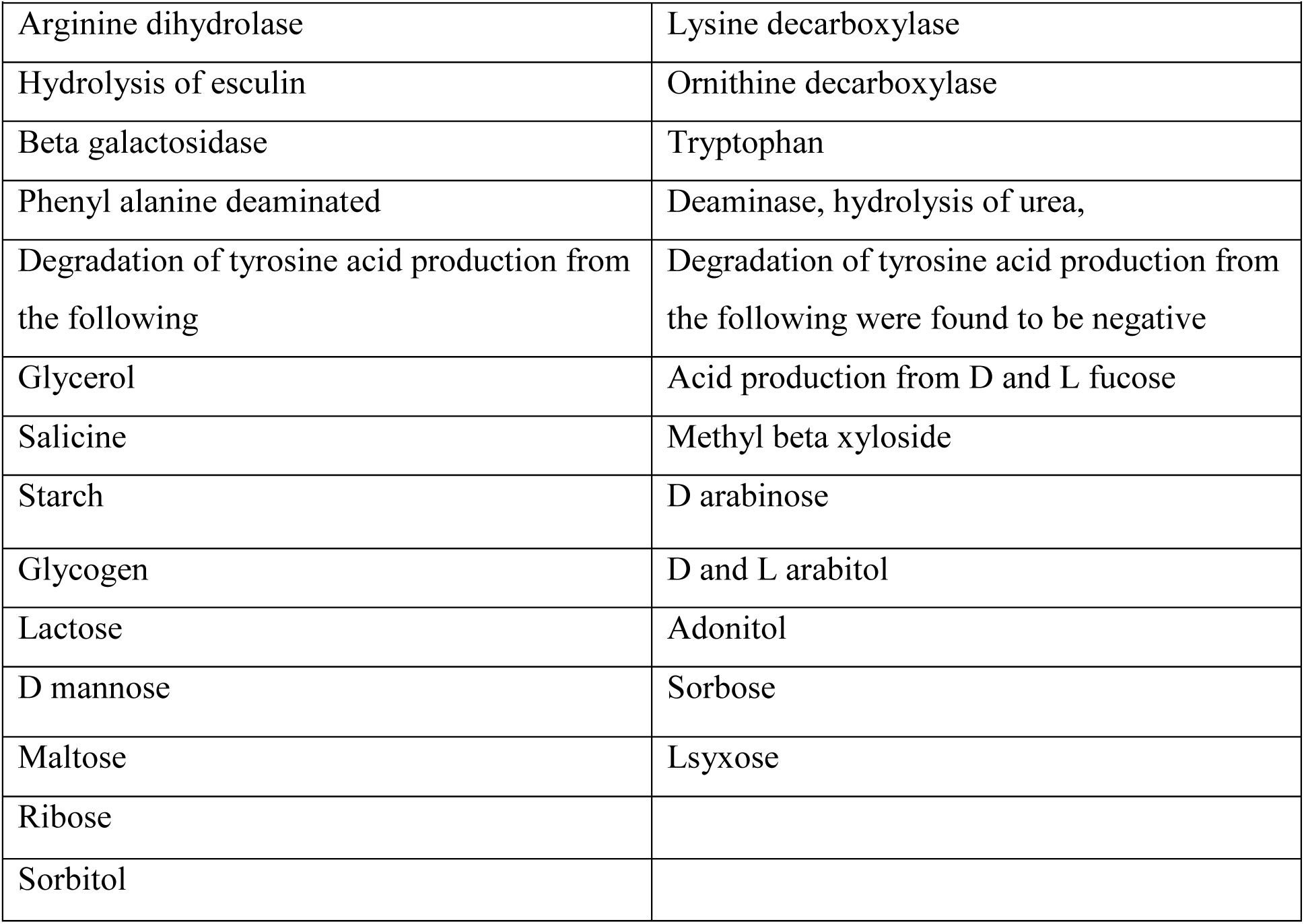
Biochemical tests of the isolated strain *Bacillus licheniformis* GPI-2

|  |  |
| --- | --- |
| **Positive tests** | **Negative tests** |
| Arginine dihydrolase | Lysine decarboxylase |
| Hydrolysis of esculin | Ornithine decarboxylase |
| Beta galactosidase | Tryptophan |
| Phenyl alanine deaminated | Deaminase, hydrolysis of urea, |
| Degradation of tyrosine acid production from the following | Degradation of tyrosine acid production from the following were found to be negative |
| Glycerol | Acid production from D and L fucose |
| Salicine | Methyl beta xyloside |
| Starch | D arabinose |
| Glycogen | D and L arabitol |
| Lactose | Adonitol |
| D mannose | Sorbose |
| Maltose | Lsyxose |
| Ribose |  |
| Sorbitol |  |

